# Conserved transcriptional co-regulation of pyrophosphate homeostasis genes governs systemic mineralization factors in mice and humans

**DOI:** 10.1101/2025.09.21.677626

**Authors:** Virgil Tamatey, Martin Várhegyi, Bence Blaha, Judith Van Wynsberghe, Dénes Juhász, Emese Bata, Dániel Tóth Márton, Muazu Muhyiddeen, Zsófia Demjén Nagy, Dániel Kovács, Áron Nagy, Anikó Ilona Nagy, Olivier Vanakker, Tamás Arányi, Flóra Szeri

## Abstract

Inorganic pyrophosphate (PPi) is a critical inhibitor of ectopic calcification, yet transcriptional regulation of genes controlling its systemic production and degradation (*ABCC6*, *ALPL*, *ANKH*, *ENPP1*) remains elusive. We hypothesized that PPi homeostasis is governed by evolutionarily conserved transcription factor (TF) network. Promoter-motif analysis of PPi genes revealed conserved enrichment of nuclear receptor TFs (ESR1, NR4A1, RXRA, NR1H3/LXRα) and metabolic regulators (SREBF1, CEBPB, HNF4A) across mouse and human orthologues. Supporting this, analysis of public RNA-seq datasets and RT-qPCR in wild-type and *Abcc6*^−/−^ mice demonstrated tight co-expression of these genes in the liver and the kidney, as central transcriptional hub of systemic PPi regulation. Functionally, analyses in mice revealed age-dependent inverse coupling between plasma PPi concentration and serum alkaline phosphatase (AP) activity, with strongest impact during early life. *Abcc6*^−/−^ mice exhibited persistently reduced although gradually increasing PPi and altered Pi/PPi ratios during aging. Translating these findings to humans, plasma PPi correlated inversely with AP activity and positively with Pi, though associations were weaker in ABCC6-deficient pseudoxanthoma elasticum patients. These results establish a conserved TF-driven program coordinating hepatic and renal expression of PPi homeostatic genes, highlight early-life sensitivity of PPi balance, and link gene-regulation to circulating mineralization factors, highlighting species-specific and pathology-driven differences.

## INTRODUCTION

Biomineralization is a tightly regulated physiological process essential for skeletal and dental development (1). It involves a highly orchestrated mechanism by which calcium phosphate-containing minerals are deposited in specific regions of the extracellular matrix (ECM), forming a scaffold for crystallization. This high spatial precision is partly achieved through the localized enzymatic removal of ubiquitous mineralization inhibitors by their respective degrading enzymes—a concept referred to as the stenciling principle (2).

A prime example of this is tissue non-specific alkaline phosphatase (TNAP) encoded by the *ALPL* gene in human and *Alpl* (*akp2*) gene in mice, which plays a pivotal role by hydrolyzing extracellular pyrophosphate (PPi) (3), a potent inhibitor of biomineralization. PPi not only regulates skeletal mineralization but also prevents hydroxyapatite crystal growth in soft tissues by binding to calcification nidi (4, 5). Under pathological conditions, ectopic calcification, the abnormal deposition of calcium phosphate in soft tissues such as arteries and cartilage, can occur, mimicking physiological mineralization (6). Notably, ectopic calcification remains a significant unmet clinical challenge: it is currently untreatable yet highly prevalent, especially with aging (7) and in common high-burden diseases such as cardiovascular and chronic kidney disease (8).

Circulatory PPi is generated from nucleotide triphosphates (NTPs), especially ATP. Most NTPs are released in a regulated manner by the ATP-binding cassette transporter ABCC6 (9), expressed predominantly in the liver and kidney, and to a lesser extent by the broadly expressed progressive ankylosis protein (homologue), ANK(H) (10). Extracellular ATP is then rapidly converted to PPi by ectonucleotide pyrophosphatase/phosphodiesterase 1 (ENPP1), the sole, ubiquitously expressed ectoenzyme capable of hydrolyzing NTPs to PPi and nucleotide monophosphates (11).

However, TNAP—either membrane-bound via its glycosylphosphatidylinositol (GPI) anchor or released into the circulation through phosphatidylinositol-specific phospholipase C (PIPLC) (12) —can degrade PPi (13), thereby promoting physiological or pathological mineralization. Circulating TNAP activity, reflected in serum alkaline phosphatase (AP) levels, is a critical determinant of bone growth and remodeling for two main reasons. First, AP activity contributes to the systemic and local inorganic phosphate (Pi) pools, essential for hydroxyapatite formation—the principal biomineral in skeletal tissues. Second, osteoblast-derived AP facilitates bone mineralization by removing the protective PPi layer from hydroxyapatite crystals, enabling further crystal growth and ECM mineralization (14).

Thus, circulating levels of Pi, PPi, and AP activity are fundamental determinants of both pathological and physiological mineralization (15), a process that is evolutionarily conserved in vertebrates.

While age is a major factor in the progression of vascular calcification, data on the kinetics of circulatory PPi levels in both humans and animal models are limited and inconsistent. Similarly, the relationship between AP activity and plasma PPi levels has not been systematically investigated. Mendelian monogenic disorders serve as valuable models for understanding the molecular mechanisms of ectopic mineralization. One such rare disease is pseudoxanthoma elasticum (PXE, OMIM 264800), a recessive multisystem disorder characterized by slowly progressive and prototypical dermatological, ocular, and vascular calcification (16). PXE typically results from biallelic fully or incompletely penetrant pathogenic variants in *ABCC6* (17, 18), though variants in both *ABCC6* and *ENPP1* can cause similar symptoms (19, 20). Due to impaired ATP release from hepatocytes, PXE patients exhibit markedly reduced circulatory PPi levels (21–23).

Since proteins involved in the same biological processes are often co-regulated at the transcriptional level via evolutionarily conserved mechanisms (24, 25), we hypothesized the existence of a genetically conserved transcriptional network that governs systemic PPi homeostasis. Specifically, we proposed that transcriptionally coordinated proteins involved in PPi production and degradation determine the systemic PPi pool supporting physiological mineralization while preventing ectopic calcification.

To explore the physiological and pathological aspects of PPi homeostasis, we first investigated whether evolutionarily conserved transcription factors have predicted binding sites in the regulatory regions of the ABCC6, ANK(H), ENPP1, and TNAP genes in both mice and humans. We then assessed the kinetics and correlations of their circulatory products (PPi, Pi, and AP activity) in wild-type and *Abcc6* / mice, as well as in human controls and PXE patients.

## MATERIALS AND METHODS

### Bioinformatics

#### Transcription factor binding site prediction

Transcription factor (TF) motifs were identified using Hypergeometric Optimization of Motif EnRichment (HOMER) v4.11 (26) and GimmeMotifs v0.24.1 (27). Analyses were based on mm10/GRCm38 (mouse) and hg38/GRCh38 (human) assemblies. Transcription start sites (TSS) for *Abcc6*, *Alpl*, *Enpp1*, and *Ank* (and human orthologs) were extracted from Ensembl v109, with promoter regions defined as –40 kb to +1 kb relative to the TSS. Motif scanning was performed in Python 3.10 using GimmeMotifs, with hits retained at p ≤ 1×10 (FPR 0.001, the default “stringent” setting,— to keep the per-site false-positive rate below 0.1 %—and has been benchmarked to give very high concordance with experimentally validated TF binding events, even when scanning large (∼10 kb) genomic regions). HOMER motif scores were converted to one-sided p-values (scoreMotifs.pl pvalue) using the same threshold. Ambiguous motif families (e.g., RXR[A/B/G]) were collapsed to a shared HGNC symbol. Only motifs detected by both tools (∼99% concordance) were considered for downstream analysis. Results were handled in pandas v2.2. TFs were considered “common” if ≥ 1 hit appeared in all four genes per species; those shared across both species were labeled “8/8 shared.”

To validate findings, promoter motif scans were repeated using the JASPAR 2024 CORE vertebrate database (28). For each gene, a –10 kb to +1 kb region (Ensembl v109 TSS) was extracted, and sequences were scanned with MEME Suite v5.5.3 (FIMO p ≤ 5×10 ; ≈ FIMO score >10 for 10-mers), using a first-order Markov background. Both DNA strands were searched; overlapping hits (same PWM, center ≥ 4 bp apart) were retained. PWM-to-gene mapping used the official JASPAR2024_vertebrates.motif2gene file. Paralogous motifs (e.g., RXRA vs RXRB) were merged if binding sites showed ≥ 80% identity (TomTom q < 0.05). For each TF, the best p-value per promoter and total significant hits were recorded. A TF was classified as “high confidence” if it met the p-value threshold (≤ 1×10 ) in ≥ 1 promoter and appeared in ≥ 3 of 4 promoters in at least one species.

#### In silico gene expression correlation analysis

We collected bulk-RNA-seq liver samples from previously published datasets (29–34). We selected GSE211975, GSE130913, GSE143206, GSE124214 and GSE174535 datasets from GEO database. Each dataset was screened manually; any treatment, genetic modification or quality failure (adapterL>L2L%, mapping rateL<L60L%) led to exclusion of the sample. In total, 63 samples remained from 8-week-old males. During raw read processing we did QCLandLtrimming using fastpL0.23.4 (automatic adapter removal, QL≥L20, readsL≥L30Lnt). We also did quality summary using multiqcL1.13; all retained runs showed mean QL≥L34 and adapter contaminationL<L1L%. For quantification, we used SalmonLv1.10.1. and GENCODE M25 (releaseL2021-03) transcriptome as index on mm10/GRCm38.p6, decoy-aware (salmon index --keepDuplicates, k-merL=L31). The gene-level summarization and normalization were executed in R 4.3.3. Filtering and grouping were performed with dplyr. For Spearman correlation and statistics, the following thresholds were used: |ρ|L≥L0.6 and BH-adjusted pL<L0.05.

#### Reagents

Unless otherwise indicated all reagents were obtained from Merck / MilliporeSigma (Burlington, Massachusetts, USA).

#### Laboratory animals

Wild-type (C57BL/6J) mice were derived from animals originally purchased from Charles River Laboratories. Abcc6L/L mice, a generous gift from prof. Arthur Bergen, were initially generated on a 129/Ola background and subsequently backcrossed to the C57BL/6J strain for more than 10 generations. (35). Mice were housed in IVC with unrestricted access to standard mouse chow (VRF1, Special Diets Services) and water according to standard housing conditions (12:12-h light:dark cycle, room temperature of 22 ± 2 °C). Genotyping biopsies also suitable for identification of the individuals were collected between postnatal days 8-14. Genotyping was performed using KAPA HotStart Mouse Genotyping Kit (Merck Life Sciences) following the manufacturer’s protocol with primers against the mouse phosphoglycerate kinase 1 promoter or Abcc6 sequence; m PGK1 reverse: ATGTGGAATGTGTGCGAGGCC; m Abcc6 forward: TGAATCTTTCTGGGGGCC AG, m Abcc6 reverse: GTACCCTGGAGCAATCCACT. Genotyping PCR reaction was performed in a T100 Thermal Cycler (Bio Rad), with the following protocol: initial denaturation: 95°C for 3 minutes; 30 cycles of denaturation: 95°C for 15 seconds, annealing: 63°C for 10 seconds; extension: 72°C for 10 seconds; final extension: 72°C for 30 seconds; Amplicons were analyzed with electrophoresis in 2% agarose gels, wild type alleles provide 163 bp, knockout 264 bp amplicons. Body weight and length (from the tip of the nose to tail base) was determined after euthanized. Experiments were carried out in accordance with the protocols approved by the Food Chain Safety and Animal Health Directorate of the Government OLce of Pest County, Hungary under permit number PE/EA/01175-6/2023. Similar number of male and female mice were used in the experiments.

#### Real-time quantitative PCR

Liver and kidney RNA was extracted using the Direct-zol RNA MiniPrep Plus kit (Zymo Research) and was used to synthesize cDNA using the RevertAid First Strand cDNA Synthesis Kit (Thermo Fisher Scientific) with oligo(dT)18 primers. Quantitative real-time PCR was performed with TB Green Premix Ex Taq II (Tli RNase H Plus (Takara) using a CFX96 Touch Real-Time PCR Detection System (Bio-Rad, Hercules, CA, USA). Primer sequences (36) are:

m Enpp1 forward ACCCTCAGTGGCAACTTGCGTT

m Enpp1 reverse TGCTTGAAGGCAGGTCCATAGC

m Abcc6 forward CACACGATGCAGCTACCAGTGA

m Abcc6 reverse GGTCATCCAGAGAAGCCTCCAT

m Ank forward CGTGGACTCATGCTGGCATTCT

m Ank reverse GTTCTCGGCATTCCAGGTGACT

m Alpl forward CCAGAAAGACACCTTGACTGTGG

m Alpl reverse TCTTGTCCGTGTCGCTCACCAT

m Rps29 forward CGGTCTGATCCGCAAATAC

m Rps29 reverse GGTCGCTTAGTCCAACTTAAT

Cycling conditions were the following: Hot start 30s 95°C and 40 cycles of 0.05s 95°C, 10s 59°C, 20s 60°C followed by a melting curve analysis from X°C to X°C. Analysis was performed using the CFX Maestro software (BioRad, Hercules, CA, USA). Relative gene expression was quantified using the ΔΔCt method with Rps29 as the reference gene.

#### Blood collection and processing in mice

To assess the kinetics and correlations of the AP-PPi/Pi axis in wild type and *Abcc6* deficient (*Abcc6*L/L ) mice we used the same phlebotomy samples to prepare plasma and sera. 500-1000 μl blood was gently drawn by cardiac puncture from isoflurane anaesthetized mice via a 23G heparinized needle and divided into Li-H300 tube (20.1309, Sarstedt) and serum-300 tubes with clotting activator (20.1308.100, Sarstedt). Blood was immediately gently mixed with the anti-coagulant or clotting activator by inverting the tubes 5 times. Tubes were centrifuged at 1000 g for 10 minutes at room temperature (RT) and plasma or sera were collected. Plasma was subsequently centrifuged again at 1000 g for 10 minutes at RT. All samples were aliquoted and stored at -80°C until further use.

#### Serum phosphate and alkaline phosphatase activity in mice

Serum concentration of inorganic phosphate (Pi) was determined using a commercial kit (MAK307-1KT, Merck Life Science) according to the manufacturer’s protocol. Serum alkaline phosphatase activity was measured in an endpoint assay in a 96-well microtiter plate. In brief, 5µl of sample was added to a total volume of 200 μL solution per well into 96-well plates (Costar 3912). The solution contained 30 mM TRIS-Glicin buffer (pH10.4), 12.5 mM MgCl2 and 3 mM 4-Nitrophenyl phosphate disodium (pNPP) salt hexahydrate (Phosphatase substrate, S0942 Merck Life Science). The plate was sealed, and the mixture was incubated for 60 minutes at RT. The absorbance at 405 nm (reflecting 4-nitrophenol production) was determined on a Enspire Multimode Plate reader (PerkinElmer). Difference in absorbance between samples and blank was calculated and alkaline phosphatase activity was quantified using calibration of serial dilutions of 4-nitrophenol (pNP) sodium salt dihydrate standard (36612 Merck Life Science) as calibration curve (0,2-12,5 mM PnP calibration).

#### Plasma inorganic pyrophosphate in mice

The PPi content of the mouse plasma was determined enzymatically, using the ATPS bioluminescent method (37), with slight modifications. Briefly, PPi was converted into ATP in an assay containing 80µM MgCl2, 50mM HEPES pH 7.4, 32mU/ml ATP Sulfurylase (M1017B, New England Biolabs, Ipswich, MA, USA) and 8µM adenosine 5’-phosphosulfate (A5508, Sigma-Aldrich, Saint Louis, MO, USA). Samples were incubated at 37°C for 30 minutes, followed by enzyme inactivation at 90°C for 10 minutes. In the next step, ATP levels were evaluated utilizing the BacTiterGlo (Promega, Madison, WI, USA) bioluminescent assay, prepared according to the manufacturer’s instructions, and analyzed in a multimode plate reader (PerkinElmer). Plasma PPi concentrations were calculated using analytical calibration standards (Biothema, Sweden) and corrected for initial plasma ATP concentrations.

### Patient cohorts

#### Zero cardiac calcification cohort

Cardiac computed tomography angiography was performed with dedicated cardiac CT scanner (CardioGraphe, GE Healthcare, Chicago, IL, USA) or a third-generation dual source CT scanner (SOMATOM Force, Siemens Healthineers, Forchheim, Germany) with ECG triggered axial acquisition mode at the Heart and Vascular Centre, Semmelweis University, Budapest, Hungary (38). Individuals with zero total cardiac calcification score, combining calcium loads of coronary arteries with that of aortic and mitral valves, were selected from prospectively enrolled patients, with exclusion criteria including bisphosphonate or vitamin K antagonist therapy, calcium, phosphate or liver metabolism disorders, diabetes, osteoporosis and impaired renal function (estimated glomerular filtration rate (eGFR) < 60 mL/min/1.73 m²). All patients gave written informed consent for CT scanning, blood sampling, and statistical data analysis. The investigation was approved by the Scientific and Research Ethics Committee of the Hungarian National Medical Scientific Council (BMEÜ/3473-1/2022/EKU). The study protocol complied with the declaration of Helsinki.

#### PXE cohort

Seventy-one PXE patients were included from the PXE Reference Center of the Center of Medical Genetics (CMGG) of the Ghent University Hospital. In all PXE patients, molecular genetic diagnosis was histologically confirmed by performing Von Kossa staining on a lesional skin biopsy to identify mid-dermal elastic fiber calcification and fragmentation. All patients gave informed consent, and the study was approved by the Ethical Committee of the Ghent University Hospital, Ghent, Belgium. The study protocol complied with the declaration of Helsinki.

#### Blood sampling and analysis in humans

Blood from patients included in the non-calcified cohort in the Heart and Vascular Centre, Semmelweis University, was drawn into serum separation vacuum tubes for the assessment of Pi and AP activity, and to K_3_EDTA vacuum tubes for plasma preparation for PPi concentration assessment. Serum samples were analyzed following the routine procedures of the Heart and Vascular Centre. The plasma fraction from the K_3_EDTA tube was isolated via two consecutive centrifugations (1000g for 5 minutes at room temperature, repeatedly) and transferred into platelet separation tubes (Vivaspin Filtrate, 300kDa, 13279-E, Sartorius, Göttingen, Germany). The platelet-free plasma was obtained by further centrifugation at 2200g for 30 min at room temperature. Samples were stored at −80°C until PPi measurement. In the Belgian PXE patient cohort blood samples were collected into vacuum citrate tubes, the plasma fraction was isolated via centrifugation (1000g for 10 minutes at at 4°C). Pi levels and AP activity were determined according to the routine procedures of the Ghent University Hospital. For PPi determination plasma was transferred into platelet separation tubes (Vivaspin Filtrate, 300kDa, 13279-E, Sartorius, Göttingen, Germany), and platelet-free plasma was obtained by further centrifugation at 2300g for 35 minutes at 4°C, and stored at −80°C until measurement.

#### Plasma inorganic pyrophosphate in humans

The PPi content of the zero calcification individuals were determined as described above for mice.

The PPi content of the PXE patients was determined similarly using the two-step ATPase luminescence assay (22). Briefly, PPi was converted into ATP in an assay containing 104µM MgCl2, 31mM HEPES pH 7.4, 825mU ATP Sulfurylase (M0394L, Bioké NV, Leiden, The Netherlands) and 64µM adenosine 5’-phosphosulfate (SC-214506, Bio-Connect, Huissen, The Netherlands). Samples were incubated at 37°C for 30 minutes, followed by enzyme inactivation at 90°C for 10 minutes. In the next step, ATP levels were evaluated utilizing the BacTiterGlo (G8230, Promega, Leiden, The Netherlands) bioluminescent assay, prepared according to the manufacturer’s instructions, and analyzed in a multimode plate reader (Glomax). Plasma PPi concentrations were calculated using an internal calibration curve and corrected for initial plasma ATP concentrations.

#### Statistical analysis and mathematical modeling

Variables were tested for normality using the Shapiro-Wilk test. Normally distributed variables are presented as mean±SEM, whereas non-normally distributed variables as median [interquartile ranges]. Variables with normal distribution are compared using unpaired two-tailed Student’s T-test, those with non-normal distribution are compared using the two-tailed Mann-Whitney U test. Pearson coefficients are reported for normally distributed variables, while Spearman’s rank correlation coefficients are used for non-normally distributed variables. Univariate and multivariate linear regressions were performed in R (glm, MASS::polr). One *Abcc6* / mouse was excluded as an outlier and altogether 30 mice due to missing values, leaving 91 *Abcc6* / and 50 wild-type mice for analysis; 66 adult PXE patients were also included. Independence and multicollinearity were assessed (Durbin–Watson test; VIF < 5), and all models met criteria. Linearity, normality, and goodness-of-fit were evaluated using residual and Q–Q plots. Statistical significance was determined at the 0.95 confidence level (p < 0.05). Analyses were performed in GraphPad Prism 10.5.0 or in R (v4.3.3).

## RESULTS

### Conservation of transcription factors and coordinated hepatic expression of PPi homeostasis genes in public RNA-seq datasets

We hypothesized that PPi homeostasis is regulated through evolutionarily conserved transcription factor–driven expression of genes controlling the systemic generation and degradation of PPi. Therefore, we first analyzed transcription factor (TF) binding sites in the regulatory regions (−10 kb/+1 kb from the TSS) of *Abcc6*, *Alpl*, *Ank*, and *Enpp1* and their human orthologues using the bioinformatics tool HOMER (26). HOMER identified several shared motifs across orthologous promoters, with NR4A1, ESR1, NR1H3/LXRa, RXRA and SREBF1 enriched in both species, while HNF4A and CEBP/B detected only in human promoters (Figure 1A). To validate these results, we applied the independent JASPAR tool (28). TF binding sites identified by both HOMER and JASPAR are marked with “#” in Figure 1A, and those unique to JASPAR with “&”. Four TFs: NR4A, ESR1, RXRA, and SREBF1, were consistently detected in both species by both tools. CEBP/B and HNF4A were reproducibly identified in human, though HOMER failed to detect HNF4A in mouse, and both tools found no CEBP/B binding motif in mice. NR1H3/LXRa was detected in both murine and human only by HOMER.

**Figure 1:**
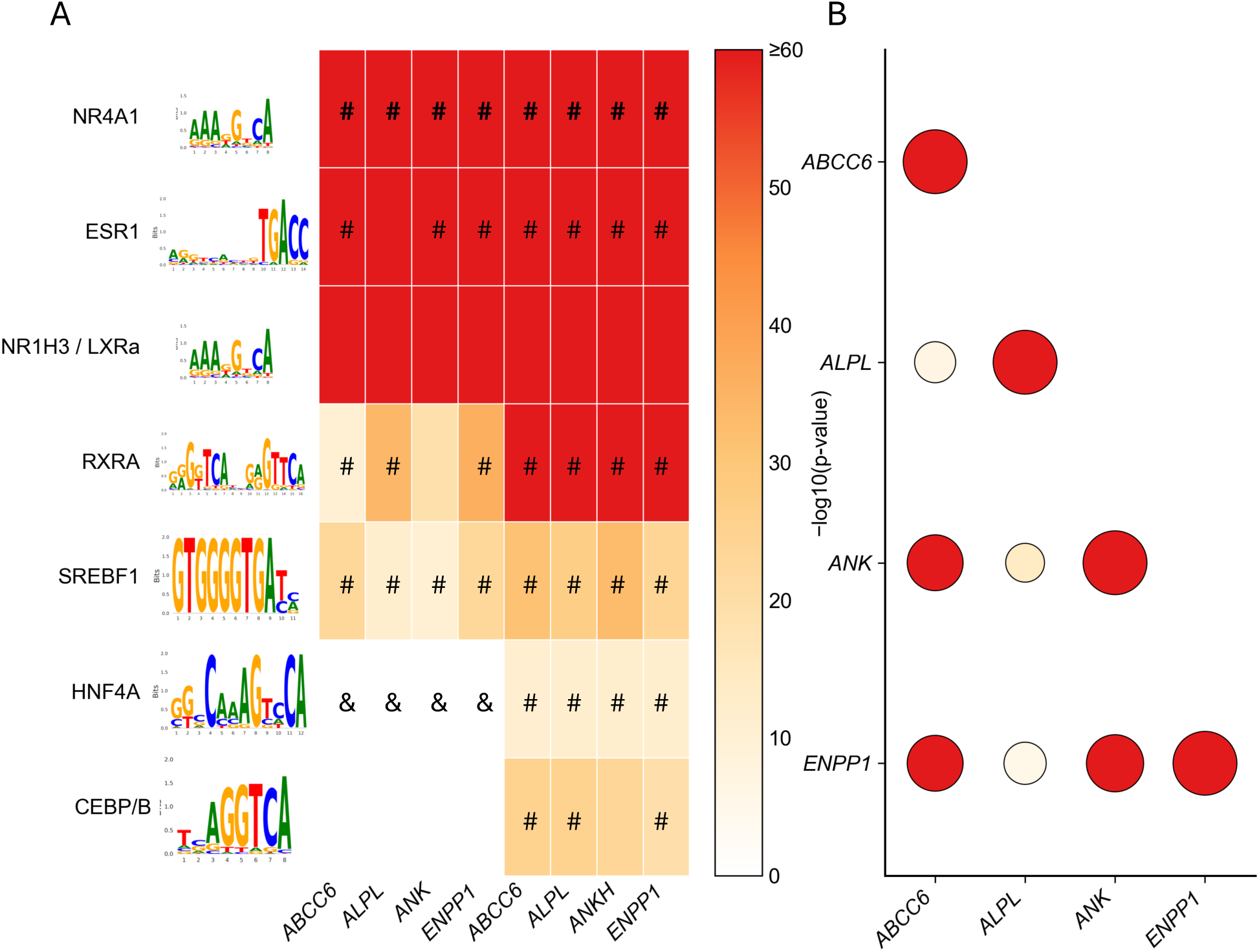
Shared transcription factor motifs and co-expression of PPi homeostatic genes. (A) Promoter motif analysis of murine and human Abcc6, Alpl, Ank, and Enpp1 using HOMER. Heatmap shows per-gene enrichment with color intensity representing −log₁₀ p-values. “#” indicates motifs supported by both HOMER and JASPAR; “&” indicates motifs identified only by JASPAR. (B) Pairwise expression correlations of PPi homeostatic genes in murine RNA-seq datasets (8-week-old males, n=63). Heatmap depicts −log₁₀ p-values by color intensity, with circle size proportional to Spearman correlation coefficient (ρ), relative to ρ = 1 along the diagonal. *Abcc6/ABCC6*, ATP binding cassette subfamily C member 6; *Alpl/ALPL*, alkaline phosphatase, liver/bone/kidney; *Ank/ANKH*, progressive ankylosis protein (homolog); CEBPB, CCAAT/enhancer-binding protein beta; *Enpp1/ENPP1*, ectonucleotide pyrophosphatase/phosphodiesterase 1; ESR1, estrogen receptor 1; HNF4A, hepatocyte nuclear factor 4 alpha; NR1H3/LXRα, nuclear receptor subfamily 1 group H member 3 (liver X receptor alpha); NR4A1, nuclear receptor subfamily 4 group A member 1; RXRA, retinoid X receptor alpha; SREBF1, sterol regulatory element binding transcription factor 1.

Together, these analyses demonstrate conserved enrichment of nuclear receptor–related factors, with NR4A1 and ESR1 showing the strongest cross-species signals, while RXRA, SREBF1, CEBP/B, and HNF4A appearing more prominent in human promoters. To validate our hypothesis on the co-regulation of the four PPi homeostasis genes we next analyzed publicly available datasets. We generated a unified dataset composed of 63 bulk RNA-seq samples from male mouse livers (29–34). In this unified murine dataset, we found high pairwise Spearman correlation (ρ > 0.80, p ≤ 3.4 × 10LL) between the expression levels of all the four PPi homeostasis genes (Figure 1B).

#### Correlation of the PPi homeostasis genes in wild-type and *Abcc6* / mice

To experimentally validate our bioinformatic findings we analyzed the gene expression of 60-day old mice by real-time quantitative polymerase chain reaction (RT-qPCR). We first investigated if the expression of the four genes (*Abcc6*, *Alpl*, *Ank* and *Enpp1*) is different in the liver and kidney of *Abcc6* wild type (wt) and *Abcc6* / (KO) littermates from *Abcc6* ^+^/ breeding pairs. We did not detect significant difference in the gene expression of the four genes of wt and KO littermates (Figure 2 A). Importantly, the disrupted *Abcc6* allele was similarly expressed to that of the wild type at the mRNA level, albeit the former transcript cannot be translated to a functional ABCC6 protein (35). As we found no genotype-specific difference in the gene expressions, we analyzed the correlations between the expression of the above genes in a pairwise manner, by plotting the expression of the wt and KO mice together. In the liver, the four PPi homeostatic genes displayed significant pairwise correlations (n = 22, Pearson r > 0.5; Figure 2B). The strongest associations were observed between Enpp1 and *Alpl* or *Ank* (r > 0.8, p < 0.0001), followed by *Ank* with *Abcc6* or *Alpl* (r > 0.7, p < 0.0001). The weakest, though still robust, correlations were between *Abcc6* and *Alpl* or *Ank* (r > 0.56, p < 0.006). Similarly, in the kidney we detected strong correlations between the four genes (n = 23, Spearman ρ > 0.7, p < 0.0001) as shown in Supplementary Figure 1.

**Figure 2:**
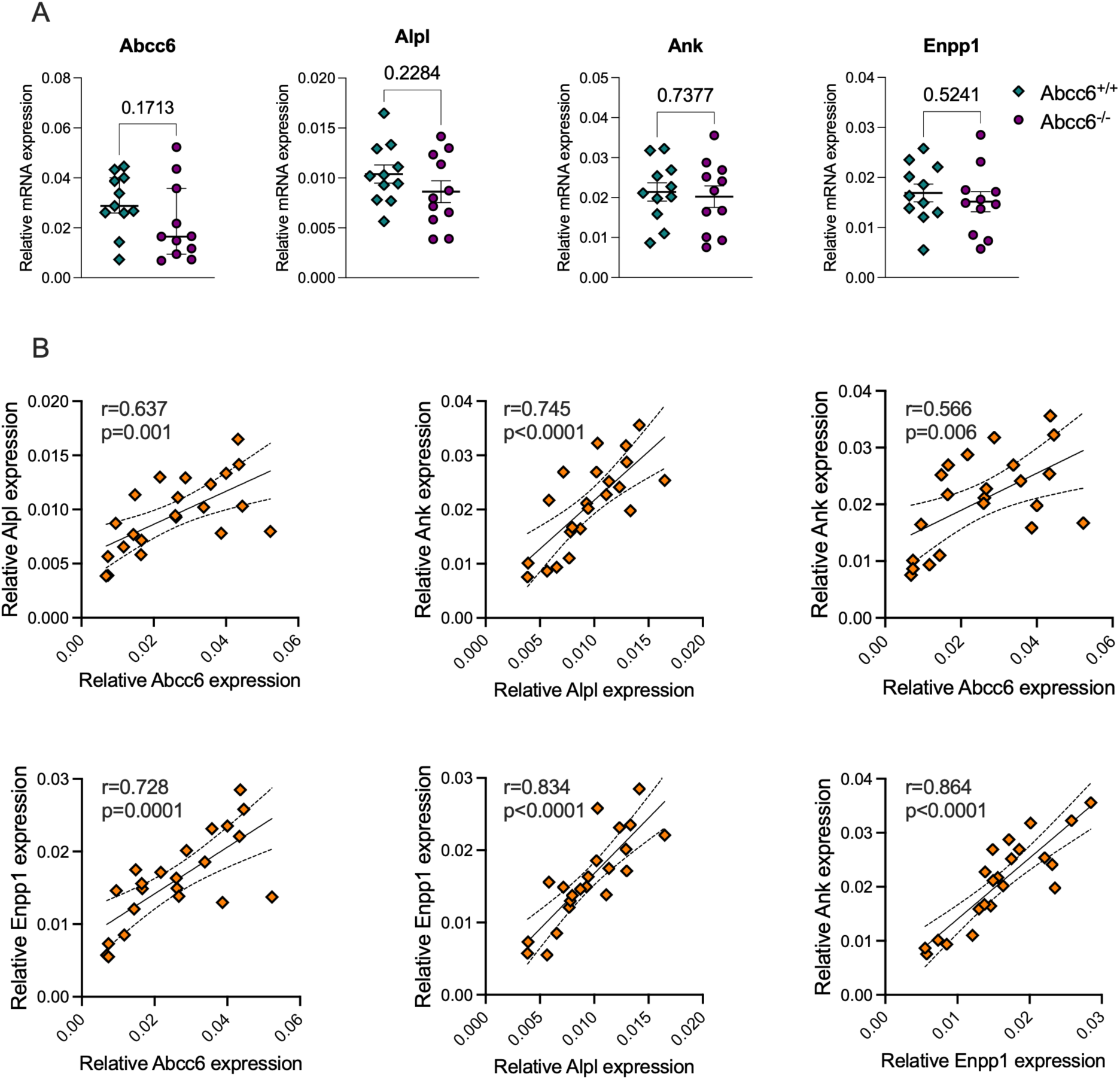
Hepatic expression and correlation of PPi homeostatic genes. (A) Expression of PPi homeostatic genes in livers of 60-day-old wild-type and Abcc6⁻/⁻ littermates (n = 11 per group), normalized to Rps29. Data are shown as mean ± SEM, except Abcc6 (median ± IQR). Statistical comparisons: Mann–Whitney U test for *Abcc6*, unpaired two-tailed Student’s t-test for other genes. (B) Hepatic expression of PPi homeostatic genes in 60-day-old littermates (n = 22). Pairwise relationships with Pearson correlation coefficients (r) and linear regression with corresponding two-tailed p-values. *Abcc6*, ATP binding cassette subfamily C member 6; *Alpl*, alkaline phosphatase, liver/bone/kidney; *Ank*, progressive ankylosis protein; *Enpp1*, ectonucleotide pyrophosphatase/phosphodiesterase.

#### Kinetics and correlation of circulatory factors in wild type and *Abcc6* / mice

Since PPi homeostatic genes exhibited significantly correlated expression in the liver, we next examined how this coordinated regulation translates into circulating mineralization products. We analyzed serum AP activity and plasma PPi concentration in wild-type and *Abcc6* / mice. Between weaning at postnatal day 21 (PND21) and 8 weeks of age, body mass increased 2.1– 2.5-fold in a near-linear manner, respectively, coinciding with bone growth reflected by a quasi-linear gain in body length that plateaued around sexual maturation (∼PND60) (Supplementary Figure 2 A). We observed a significant decline in serum AP activity from weaning to 8 weeks of age in both wild-type and *Abcc6* / mice, while plasma PPi levels increased substantially (∼1.5–2-fold, respectively) (Figure 3 A), along with body weight (Supplementary Figure 2 A). Although AP activity did not differ between genotypes with age (Supplementary Figure 2 B), plasma PPi in *Abcc6* / mice remained approximately half of wild-type levels throughout, consistent with the absence of functional ABCC6 (Supplementary Figure 2 C). Nevertheless, remarkably, PPi in *Abcc6* / mice increased twofold by 8 weeks, similar to the >1.5-fold rise in wild-type mice (Figure 3 A). No sex-specific differences were detected (Supplementary Figure 2 B, C).

**Figure 3:**
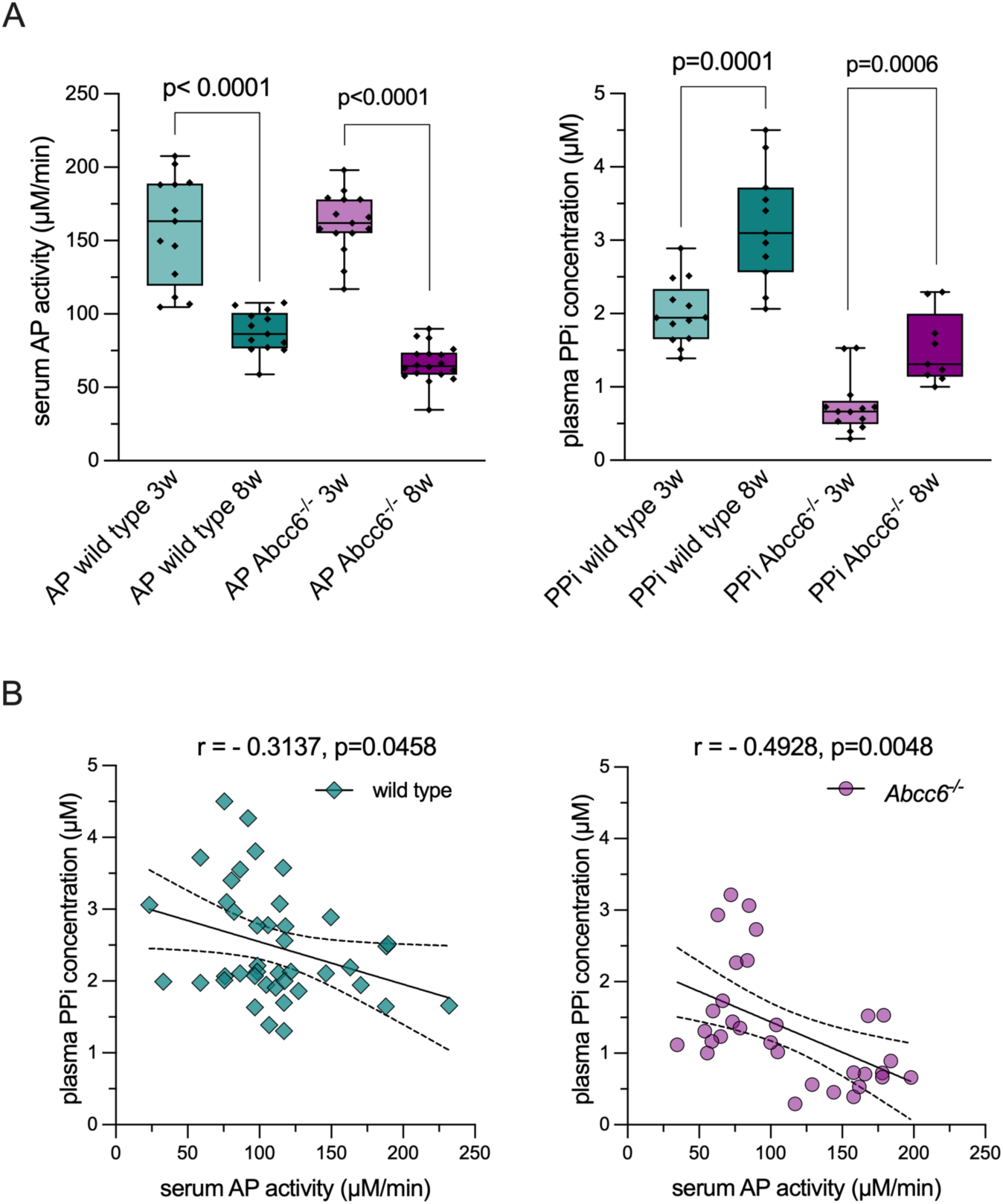
Relationships between alkaline phosphatase activity and plasma PPi concentration in young wild-type and *Abcc6*⁻/⁻ mice. (A) Serum AP activity and plasma PPi levels in 3- to 8-week-old wild-type and *Abcc6*⁻/⁻ mice. Box plots show medians with interquartile ranges, minimum and maximum values, and individual data points. Significancy was tested using Mann– Whitney U test (p < 0.05). (B) Correlation of plasma PPi concentration with AP activity in young wild-type (n = 41) and *Abcc6*⁻/⁻ (n = 31) mice between postnatal day (PND) 21 and PND 62. Linear regression with 95% confidence intervals is shown. Spearman correlation coefficients (ρ) and two-tailed p-values are indicated. Teal diamonds represent wild-type and purple circles *Abcc6*⁻/⁻ mice, with transparency used to highlight overlapping data points. AP, alkaline phosphatase; PPi, pyrophosphate.

Given the concerted expression of PPi homeostatic genes, we investigated the correlations between circulatory mineralization factors, and found that in young *Abcc6* / mice, PPi levels strongly inversely correlate with AP activity (Spearman ρ ≈ -0.5, p = 0.005, n = 31), whereas wild-type mice show a weaker, yet significant, negative correlation (ρ ≈ -0.3, p = 0.045, n = 41) (Figure 3 B). Extending the analysis to adult wild-type mice (up to PND200, n = 72), serum AP activity declines sharply during maturation but remains stable thereafter (Figure 4A). A two-phase exponential model fit AP activity, with strong negative correlation with age (ρ ≈ -0.85, p < 0.0001). Plasma PPi and the Pi/PPi ratio remains unchanged with age, while AP activity shows a weak positive correlation with Pi/PPi (ρ ≈ 0.25, p = 0.040) but with PPi (Figure 4B). Univariate linear regression analysis identifies age as the only significant, negative contributor to the Pi/PPi ratio (p = 0.029), but this effect is lost in the multivariate linear regression model including sex and serum AP activity.

**Figure 4:**
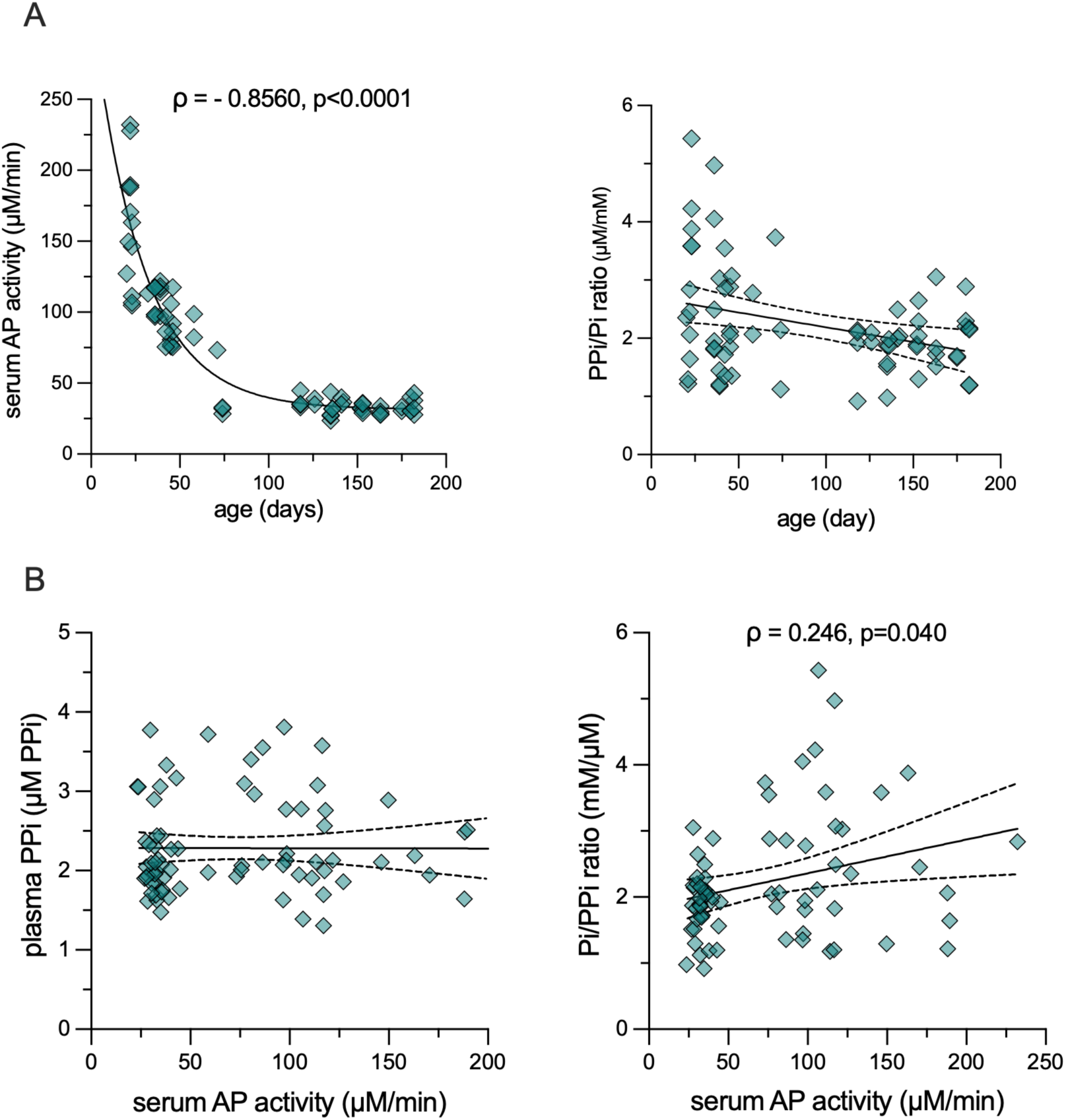
Relationships between age, AP activity, PPi and Pi/PPi ratio in wild-type mice. (A) Age dependence of serum alkaline phosphatase (AP) activity and Pi/PPi ratio. (B) Correlation between serum AP activity and PPi or the Pi/PPi ratio. Linear regression with 95% confidence intervals is shown. Spearman correlation coefficients (ρ) and corresponding two-tailed p-values are indicated at significant correlations. Teal diamonds represent individual wild-type mice (n = 77), with transparency used to highlight overlapping data points. AP, alkaline phosphatase; PPi, pyrophosphate; Pi/PPi, serum phosphate/ plasma pyrophosphate ratio.

In *Abcc6* / mice up to 200 days, AP activity declined markedly with age, however, unlike wild-type mice, PPi levels positively, while the Pi/PPi ratio negatively correlated with age. Extending the analysis to 600 days (n = 101) confirmed that AP activity fits a two-phase exponential curve (ρ ≈ -0.83, p < 0.0001) (Figure 5A) alike wild type. However, unlike the wild type mice, *Abcc6* / mice show a persistent age-dependent increase in PPi (ρ ≈ 0.54, p < 0.0001) and a robust decline in the Pi/PPi ratio (ρ ≈ -0.6, p < 0.0001) throughout the lifespan (Figure 5 A). Furthermore, AP activity correlates inversely with PPi (ρ ≈ -0.4, p < 0.0001) and positively with the PPi/Pi ratio (ρ ≈ 0.39, p < 0.0001) (Figure 5B). No sex-specific differences are apparent (Supplementary Figure 3). Univariate linear regression models reveal an inverse correlation of the Pi/PPi ratio with age (p < 0.0001) and a positive correlation with AP activity (p < 0.0001). In multivariate linear regression adjusted for age and sex (p = 1.3 × 10 , adjusted R² = 0.305), AP activity remains a significant predictor of the Pi/PPi ratio (p < 0.0001) explaining over 30% of variance in Pi/PPi ratio, whereas age no longer contributes.

**Figure 5:**
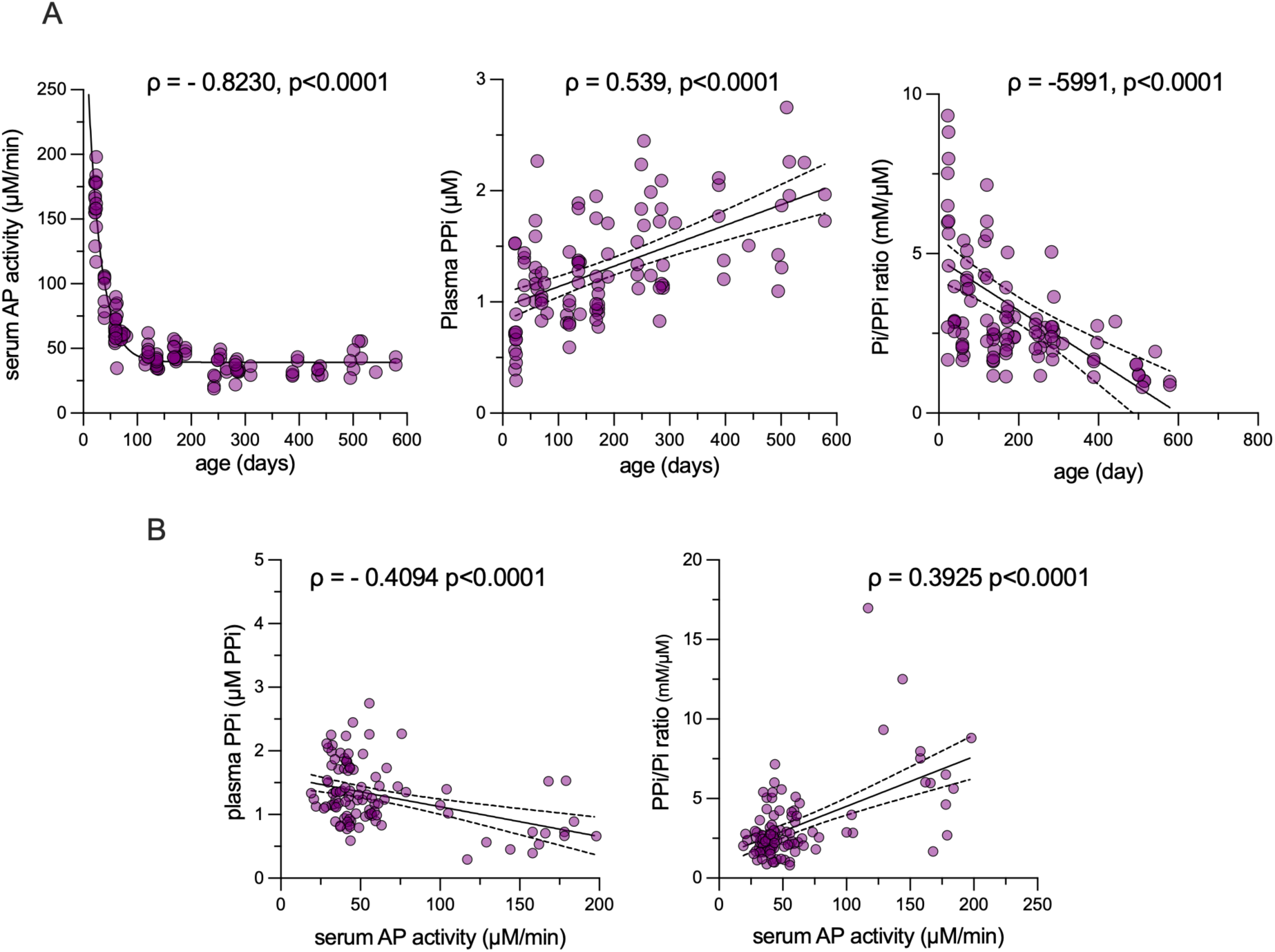
Relationships between age, AP activity, PPi, and Pi/PPi ratio in *Abcc6^-/-^* mice. (A) Age dependence of serum alkaline phosphatase (AP) activity, PPi concentration and Pi/PPi ratio. (B) Correlation between serum AP activity and PPi or the Pi/PPi ratio. Linear regression with 95% confidence intervals is shown. Spearman correlation coefficients (ρ) and corresponding two-tailed p-values are indicated at significant correlations. Purple circles represent individual *Abcc6^-/-^* mice (n = 101), with transparency used to highlight overlapping data points. AP, alkaline phosphatase; PPi, pyrophosphate; Pi/PPi, serum phosphate/ plasma pyrophosphate ratio.

#### Correlation between human circulatory factors

Next, we investigated the factors of the PPi homeostasis in humans. To generate a cohort with individuals negative for ectopic calcification, we prospectively recruited patients referred for cardiac CT and included them into our study if they had zero cardiac calcification. In total we included 26 patients, including 18 males, with a broad age distribution (median age = 55 years; IQ range 49.5, 55.25 years). In these patients, we found no significant correlation between serum AP activity, plasma PPi or serum Pi levels with age. However, plasma PPi concentration exhibited a strong inverse correlation with AP activity (Spearman ρ ≈ -0.47, p=0.014), while the Pi/PPi ratio positively correlated with AP activity (Pearson r ≈ 0.45, p=0.020). In addition, in these individuals the plasma PPi level positively correlated with serum Pi concentration (Spearman ρ ≈ 0.61, p = 0.0008) (Figure 6. A). Given the small cohort size (n = 26), no statistical models were generated.

**Figure 6:**
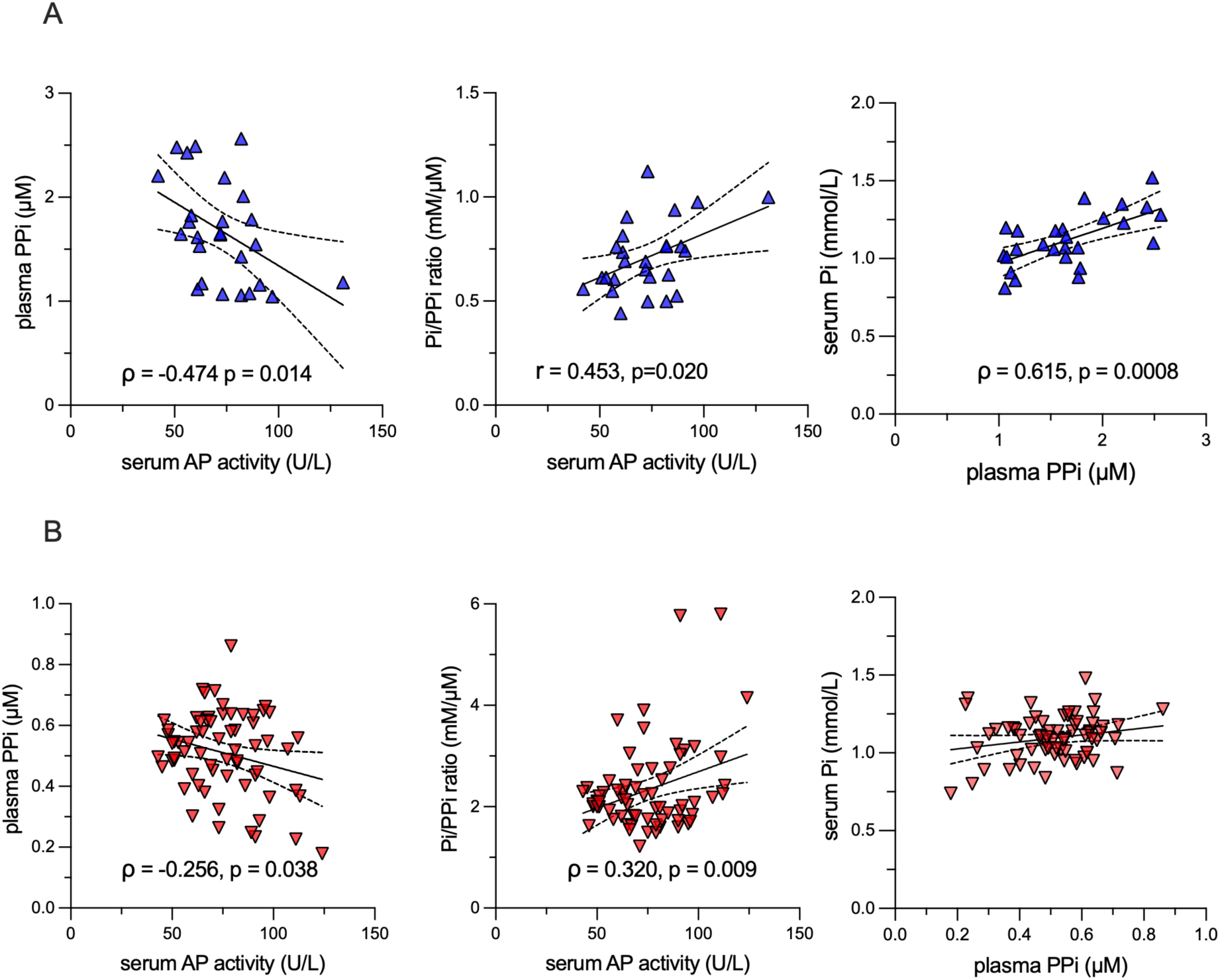
Relationships between AP activity, PPi, and Pi/PPi ratio in human. (A) Correlation between serum AP activity and PPi or the Pi/PPi ratio, or PPi and Pi levels in adult patients without detectable cardiac calcification (n = 26). (B) Correlation between serum AP activity and PPi or the Pi/PPi ratio, or PPi and Pi levels in adult patients without detectable cardiac calcification (n = 66). Blue upward triangles represent individual patients without detectable cardiac calcification; red downward triangles represent PXE patients. Transparency of symbols reveals overlapping data. Linear regression with 95% confidence intervals is shown. Spearman or Pearson correlation coefficients (ρ or r) and corresponding two-tailed p-values are indicated at significant correlations. AP, serum alkaline phosphatase; PPi, pyrophosphate; Pi, serum phosphate; Pi/PPi, serum phosphate/ plasma pyrophosphate ratio.

We were also interested in the above correlations in ABCC6 deficient patients. The Belgian pseudoxanthoma elasticum cohort comprised of 71 individuals including five children, whom, we excluded them from the analysis. The study cohort comprising 66 adult PXE patients including 25 males had a median age of 47.5 years (IQ range 32.5, 54.25 years). Similarly to the patient cohort with zero cardiac calcification, in the PXE cohort no significant correlations were found between age and AP activity, PPi levels, or Pi levels. Furthermore, we only found a significant week negative association between AP activity and PPi levels (Pearson r ≈ -0.26, p = 0.038) and a somewhat stronger positive correlation between AP activity and Pi/PPi levels (Pearson r = 0.32, p = 0.009) (Figure 6. panel B). In PXE patients, univariate analysis identified AP activity as a significant contributor to the Pi/PPi ratio (p = 0.009), unlike age or sex. The association of the Pi/PPi ratio with AP activity remained significant (p = 0.018) in multivariate linear regression adjusted for age and sex. Although the model explained a modest proportion of variance (adjusted R² = 0.077), this finding highlights AP activity as an independent factor influencing the Pi/PPi ratio in PXE.

## DISCUSSION

### Conserved transcriptional regulation of PPi homeostasis genes

Genes of key physiological pathways are often co-regulated at the transcriptional level, through shared TF binding. We hypothesized, that PPi homeostasis genes are co-regulated hence may share evolutionary conserved TF binding sites. Our motif analysis revealed conserved enrichment of nuclear receptor TFs in murine and human PPi homeostasis genes. ESR1, NR4A1, RXRA, and SREBF1 were consistently identified across species and tools, implicating them as evolutionarily conserved regulators of *Abcc6*/*ABCC6*, *Alpl*/*ALPL, Ank*/*ANKH*, and *Enpp1*/*ENPP1*. By contrast, CEBPB was absent and HNF4A only sporadically detected in mouse promoters, although both were robustly identified in humans, consistent with prior in-vitro studies (39–41). Notably, experimental evidence supports HNF4A-mediated regulation of the murine *Abcc6* promoter (42). Yet, this divergence in CEBPB and possibly in HNF4A sites may reflect species-specific differences in transcriptional control. Overall, our findings are consistent with the established role of nuclear receptors in metabolic regulation, as well as with previous reports linking ESR1 and RXRA to vascular calcification (43, 44) and bone metabolism (45) and suggest a partially conserved, orchestrated regulation of PPi homeostasis genes.

The consistent co-expression of PPi homeostatic genes across multiple RNA-seq datasets, further confirmed by RT-qPCR in both wild-type and Abcc6 / mice, supports the concept that hepatic and renal transcriptional programs may act in concert to regulate systemic PPi availability. This regulation aligns with the previously established central role of the liver in systemic PPi homeostasis (9), with our findings supporting the liver and potentially the kidney as prime transcriptional hubs for PPi regulation. In skeletal tissues a similar phenomenon was observed in knock-out murine models revealing a concerted regulation of PPi and osteopontin by *Alpl*, *Enpp1*, and *Ank* (46). These observations underscore the possibility that shared TF networks ensure coordinated transcriptional control of PPi metabolism across tissues and species.

### Age-dependent coordination of AP activity and PPi levels in mice

Plasma PPi kinetics have not been investigated in young mice, although TNAP is known to be highly expressed in early life, resulting in elevated AP activity (47, 48). Given that PPi is a direct TNAP substrate, we hypothesized that plasma PPi availability may be influenced during this developmental window, a possibility not previously addressed. Supporting this rationale, in lambs we previously reported low postnatal PPi levels that rose sharply to reach adult concentrations by weaning (49). In our murine models AP activity exhibited a biphasic decline with age, stabilizing in adults, in line with data from 90–135-day old mice (50), while plasma PPi levels also plateaued. These dynamics are consistent with TNAP-mediated hydrolysis of PPi. Another novel finding of the present work is the strong reciprocal relationship between plasma PPi concentration and serum AP activity during post-weaning maturation, with a marked decline in AP activity paralleled by a doubling of plasma PPi levels. The disappearing strength of this inverse correlation in adulthood suggests that additional mechanisms may also shape PPi homeostasis beyond. Our work suggest that developmental regulation of AP activity critically shapes extracellular PPi availability in early life. Lambs showing a similar postnatal increase in PPi raises the possibility that such dynamics represent a conserved developmental program, which support our notion on the concerted transcriptional regulation of the PPi homeostatic genes. Taking in consideration the potency of prenatally supplemented PPi to halt ectopic calcification in Enpp1 KO offspring (37), our results highlight early age as a critical therapeutic window to combat later-in-life-manifesting ectopic calcification.

### Divergent regulation in wild-type and *Abcc6* / mice

Although wild-type and Abcc6 / animals followed broadly similar developmental trends, the knockout mice consistently showed ∼50% lower plasma PPi and, importantly, a distinct pattern of regulation. In wild-type mice, univariate regression identified age as the only significant contributor to the Pi/PPi ratio, but this effect was lost after inclusion of sex and serum AP activity in a multivariate model. This pattern suggests that the apparent influence of age on the Pi/PPi ratio in wild-type animals is at least partly explained by covariation with AP activity (and/or sex), rather than reflecting an independent effect of age per se.

By contrast, Abcc6 / mice show a clearer, more robust regulatory signal. Univariate analyses revealed a strong inverse relationship between Pi/PPi and age and a strong positive relationship between Pi/PPi and AP activity. In a multivariate model adjusted for age and sex the model remained significant and explained over 30% of the variance in Pi/PPi; serum AP activity remained an independent, strong predictor while age no longer contributed. Thus, in the Abcc6 / background AP activity accounts for a substantial fraction of the variation in the Pi/PPi balance, whereas the effect of age appears to be mediated through or confounded by AP activity. Biologically, these results are consistent with partial compensation for loss of ABCC6 (for example via Ank or other PPi-regulating pathways) and indicate that AP activity is a key modulator of the Pi/PPi ratio in the knockout animals. Finally, whereas the multivariate model for Abcc6 / mice achieves significant explanatory power, the much weaker adjusted R² in wild-type mice implies that additional, unmeasured processes contribute to Pi/PPi regulation in the intact background, an important caveat and an avenue for future mechanistic studies.

### Human validation and translational relevance

In humans, plasma PPi levels inversely correlated with AP activity, consistent with our murine findings. In individuals with zero cardiac calcification, PPi also correlated positively with serum Pi, while AP activity directly associated with the Pi/PPi ratio, a clinically relevant marker of calcification risk. This highlights the tight coupling of PPi and Pi metabolism and reinforce AP as a central physiological regulator, while raising the possibility that coordinated transcriptional control of PPi homeostasis genes contributes to this balance, a hypothesis warranting future investigation. In the disease setting, Pi/PPi ratio was recently shown to be a significant determinant of the aortic valve calcification in cardiovascular patients (38), also exhibiting a negative association between PPi and AP activity, similarly to patients with chronic kidney disease (51), in which PPi not only showed an inverse correlation with AP activity but a positive correlation with Pi levels. An earlier study however found a weak baseline correlation of plasma PPi with age and serum phosphate in CKD (52). In our PXE patient cohort the correlations were weaker compared to the zero calcification cohort, with only a modest inverse association between AP activity and PPi levels. Nevertheless, univariate regression identified AP activity as a significant contributor to the Pi/PPi ratio which remained significant in a multivariate model adjusted for age and sex. Although the model explained only a modest proportion of variance, these findings demonstrate that AP activity is an independent determinant of the Pi/PPi ratio in PXE, underscoring its role as a key modifier of extracellular PPi availability even in the disease setting. A striking difference emerged between Abcc6-deficient mice and our PXE patients: while PPi levels increased progressively with aging in knockout mice in contrast with wild-type mice, it remained stable with age in our PXE cohort. Of note, previously, in a large PXE cohort PPi levels showed a positive correlation with age (23), similarly to heterozygous carriers and PXE patients but controls (22). The controversial data warrant further investigation. However, identifying the mechanisms driving the strong ABCC6 dependent increase in PPi levels with age in mice may uncover therapeutic targets to modulate disease onset and progression in humans.

### Limitations

Several limitations warrant consideration. Our promoter motif analyses rely on computational predictions and require functional validation (e.g., ChIP-seq, reporter assays) to confirm TF binding and regulatory activity. The correlations observed in mice, although robust, do not establish causality, and further mechanistic studies are needed to define how TF networks and AP activity integrate to regulate systemic PPi. Finally, our human cohorts, while well-characterized, were relatively modest in size. Also, differences in preanalytical and analytical procedures between the zero-calcification and PXE cohorts in the two centers may have contributed, at least in part, to the observed phenomena. Larger, longitudinal studies with harmonized protocols are required to validate the observed relationships in the general population and in patient cohorts. Nonetheless, the consistency of associations across models and cohorts supports the robustness of our main conclusions.

### Implications and future directions

Our findings highlight three key implications. First, PPi homeostasis is under coordinated transcriptional control, with conserved nuclear receptor and metabolic TFs acting as candidate regulators. This opens opportunities for therapeutic modulation of PPi levels via transcription factor–targeted interventions (e.g., ESR1 or RXRA ligands). Second, developmental regulation of AP activity emerges as a critical determinant of PPi availability, suggesting that early-life may be particularly vulnerable to disruptions in PPi homeostasis, which therefore may serve as a potential therapeutic window. Third, the correlations we observed between PPi, AP activity, and the Pi/PPi ratio in humans underscore the translational relevance of murine models. Further studies dissecting the TF regulatory landscape and the mechanisms underlying the age-related compensatory increase in PPi levels in mice may reveal additional, and potentially druggable targets to combat ectopic calcification.

## Conclusion

In summary, our integrative analyses reveal that PPi homeostasis is transcriptionally co-regulated by a conserved transcription factor network, coordinated in the liver and the kidney. Furthermore, PPi homeostasis is dynamically shaped by age-dependent changes in AP activity. The strong reciprocal relationship between AP activity and plasma PPi observed in both mice and humans highlights the conserved mechanism of extracellular PPi regulation. These findings not only advance our understanding of the molecular underpinnings of systemic PPi homeostasis but also provide a framework for exploring therapeutic strategies to modulate PPi in calcification disorders such as PXE.

## Abbreviations

Abcc6/ABCC6: ATP Binding Cassette Subfamily C Member 6
ADP: Adenosine Diphosphate
Alpl/ALPL: Alkaline Phosphatase, Liver/Bone/Kidney
AMP: Adenosine Monophosphate
Ank/ANKH: Progressive Ankylosis Protein (Homologue)
AP: Alkaline Phosphatase
ATP: Adenosine Triphosphate
CEBPB: CCAAT/Enhancer-Binding Protein Beta
ECM: Extracellular Matrix
Enpp1/ENPP1: Ectonucleotide Pyrophosphatase/Phosphodiesterase 1
ESR1: Estrogen Receptor 1
GPI: Glycosylphosphatidylinositol
HNF4A: Hepatocyte Nuclear Factor 4 Alpha
NTP: Nucleotide Triphosphates
NR1H3/LXRa: Nuclear Receptor Subfamily 1 Group H Member 3 (Liver X Receptor Alpha)
NR4A1: Nuclear Receptor Subfamily 4 Group A Member 1
PGK1: Phosphoglycerate Kinase 1 Pi Inorganic Phosphate
PIPLC: Phosphatidylinositol-Specific Phospholipase C
PPi: Inorganic Pyrophosphate
PXE: Pseudoxanthoma Elasticum
RT-qPCR: Real-Time Quantitative Polymerase Chain Reaction
RpS29: Ribosomal Protein S29
RXRA: Retinoid X Receptor Alpha
SREBF1: Sterol Regulatory Element Binding Transcription Factor 1
TNAP: Tissue Non-Specific Alkaline Phosphatase
WT: Wild Type

---

**Supplementary Figure 1:**
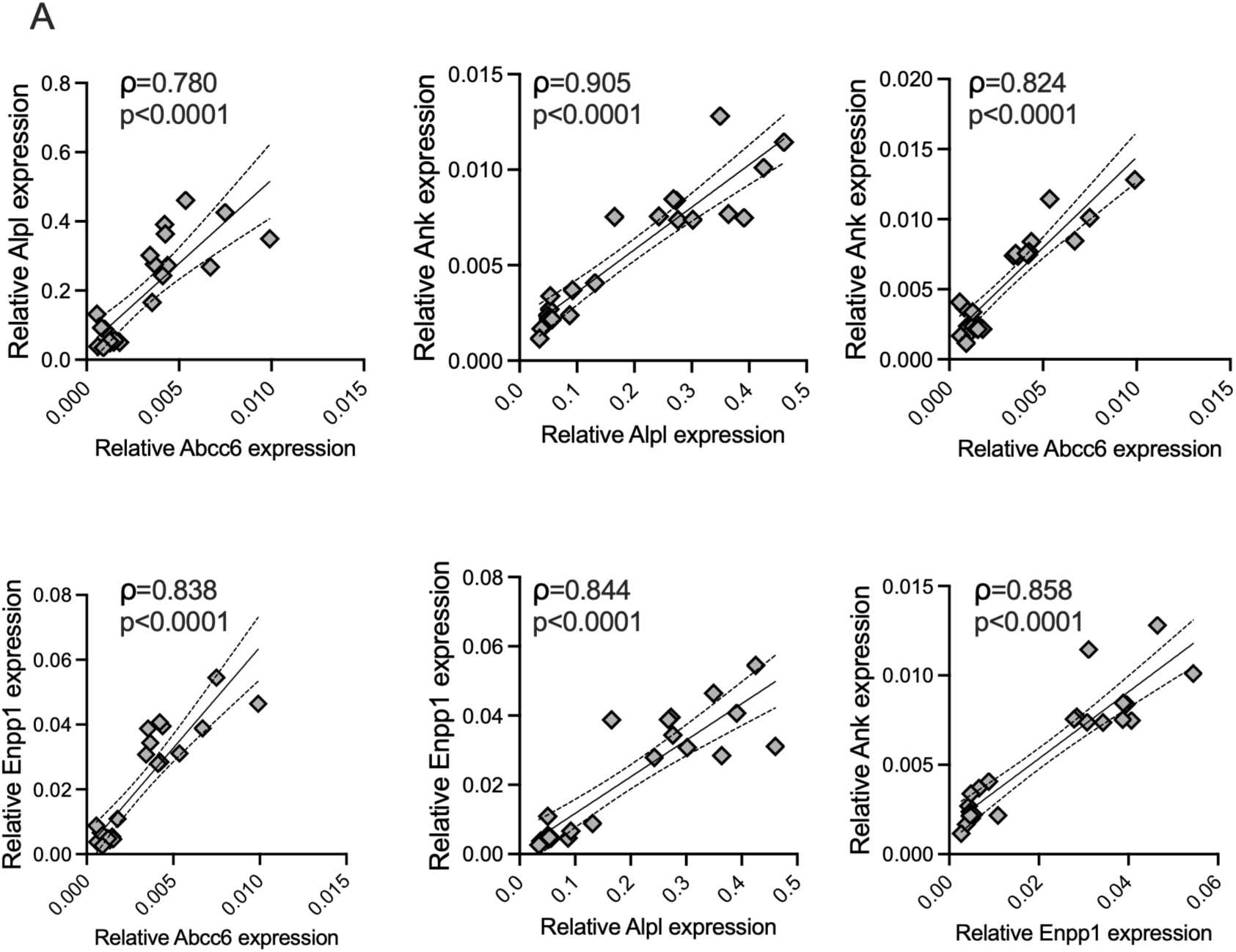
Correlation of PPi homeostatic genes in mouse kidney. (A) Pairwise correlations of the genes in kidneys of 60-day-old wild-type and Abcc6⁻/⁻ littermates (n = 23) normalized to Rps29. The Spearman correlation coefficients (ρ) and linear regression with corresponding two-tailed p-values are indicated. *Abcc6*, ATP binding cassette subfamily C member 6; *Alpl*, alkaline phosphatase, liver/bone/kidney; *Ank*, progressive ankylosis protein; *Enpp1*, ectonucleotide pyrophosphatase/phosphodiesterase.

**Supplementary Figure 2:**
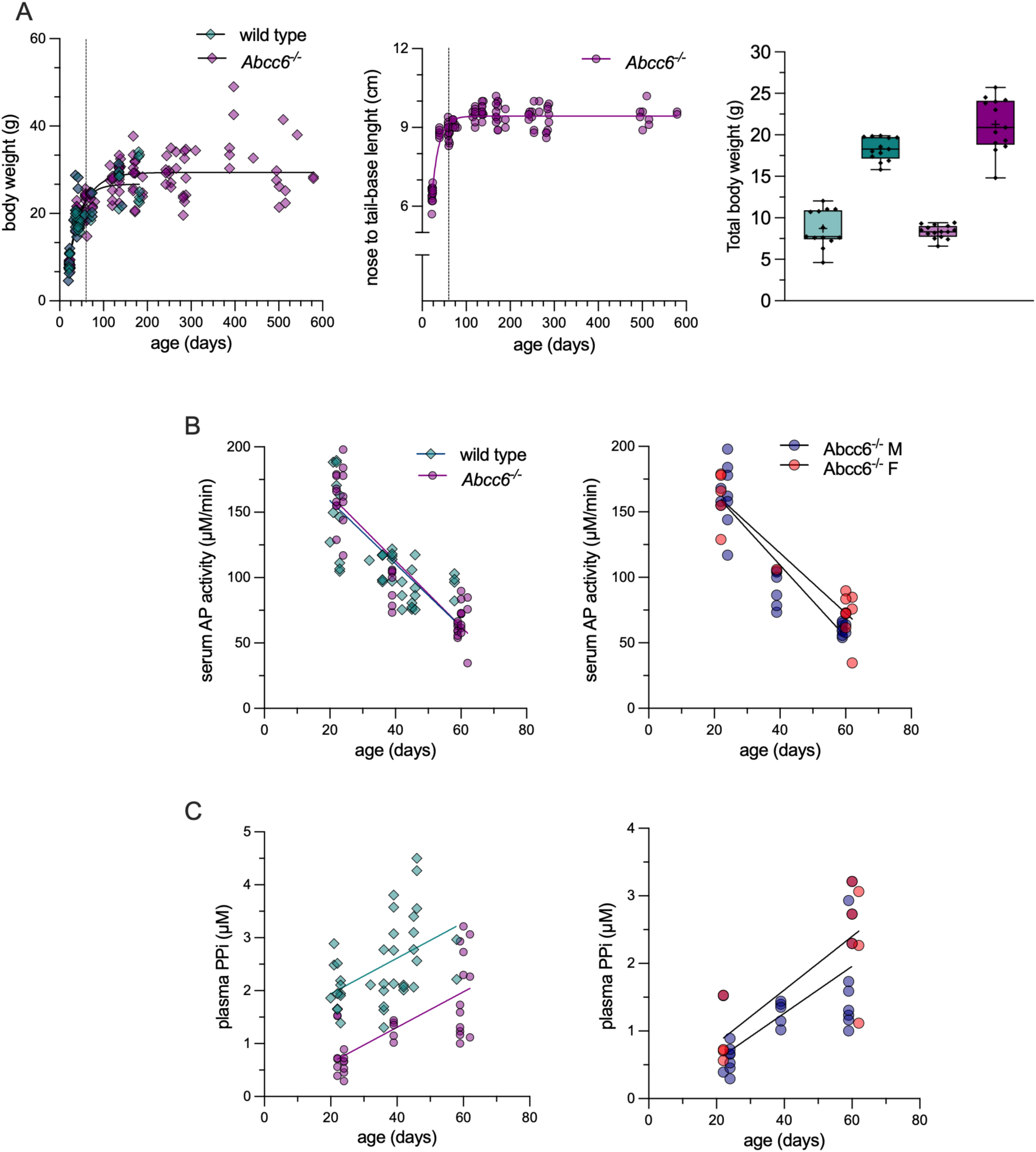
(A) Age dependence of body weight and lenght with fitted curve and the body weight of wild-type and Abcc6⁻/⁻ mice at 3nd 8th weeks of age. Box plots show medians with interquartile ranges, minimum and maximum values, and individual data points. (B) Serum AP activity as a function of age in wild-type and Abcc6⁻/⁻ mice, or in male and female Abcc6⁻/⁻ mice with linear regression. (C) Plasma PPi concentration as a function of age in wild-type and Abcc6⁻/⁻ mice, or in male and female Abcc6⁻/⁻ mice, with linear regression. Teal represents individual wild-type and purple Abcc6⁻/⁻ mice, or blue and red circles represent individual male and female *Abcc6^-/-^* mice. Transparency is used to highlight overlapping data points, when applicable. AP, serum alkaline phosphatase; PPi, plasma pyrophosphate.

**Supplementary Figure 3:**
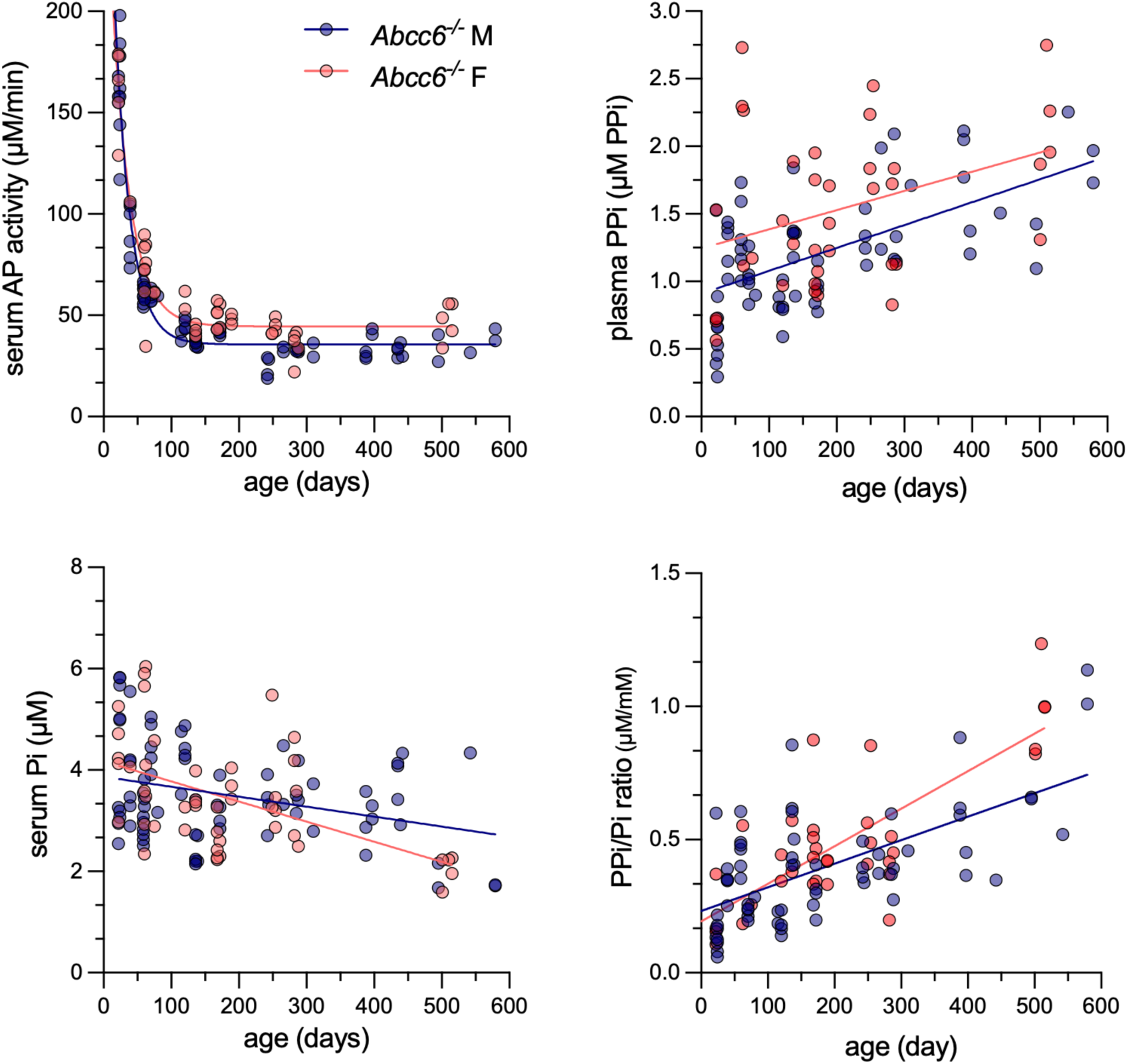
(A) Age dependence of AP activity, PPi and Pi concentration, and the Pi/PPi ratio in male and female Abcc6⁻/⁻ mice. (B) Relationsip between AP activity, PPi and Pi concentration, and the Pi/PPi ratio in male and female Abcc6⁻/⁻ mice. Blue and red circles represent individual male and female *Abcc6^-/-^* mice, with transparency used to highlight overlapping data points. Linear regression lines are indicated. AP, serum alkaline phosphatase; PPi, pyrophosphate; Pi, serum phosphate; Pi/PPi, serum phosphate/ plasma pyrophosphate ratio; M, male; F, female

